# Incorporation of Heterogeneity in a Case-Control Study Through a Mixture Model

**DOI:** 10.1101/2020.09.09.290437

**Authors:** Subrata Paul, Stephanie A. Santorico

## Abstract

Most common human diseases and complex traits are etiologically heterogeneous. Genome-wide Association Studies (GWAS) aim to discover common genetic variants that are associated with complex traits, typically without considering heterogeneity. Heterogeneity, as well as imprecise phenotyping, significantly reduces the power to find genetic variants associated with human diseases and complex traits. Disease subtyping through unsupervised clustering techniques such as latent class analysis can explain some of the heterogeneity; however, subtyping methods do not typically incorporate heterogeneity into the association framework. Here, we use a finite mixture model with logistic regression to incorporate heterogeneity into the association testing framework for a case-control study. In the proposed method, the disease outcome is modeled as a mixture of two binomial distributions. One of the component distributions refers to the subgroup of the population for which the genetic variant is not associated with the disease outcome and another component distribution corresponds to the subgroup for which the genetic variant is associated with the disease outcome. The mixing parameter corresponds to the proportion of the population for which the genetic variant is associated with the disease outcome. A simulation study of a trait with differing levels of prevalence, SNP minor allele frequency, and odds ratio was performed, and effect size estimates compared between the models with and without incorporating heterogeneity. The proposed mixture model yields lower bias of odds ratios while having comparable power compared to classical logistic regression.

## Introduction

Most human diseases, such as diabetes, cardiovascular disease, and cancer, are complex traits. Complex traits, being the result of many genetic variants, their interactions, and interaction with environmental factors, are heterogeneous with respect to disease pathophysiology; however, it is not generally easy to determine the heterogeneity based on phenotypic symptoms. To illustrate how heterogeneity affects an association study, suppose, in a case-control study we want to test whether a SNP, S, is associated with a disease trait, T, using a sample of 1000 cases and 50,000 controls. Assume that the disease has two underlying subtypes with 100 cases corresponding to type 1 and the remainder corresponding to type 2. If the SNP S is highly associated with disease type 1 but not with type 2, we are likely to be underpowered to detect an association based on the heterogeneous mixture of cases. In such situations the effect sizes are underestimated in magnitude. This simple example of heterogeneity is similar to phenotypic misclassification. Phenotypic misclassification can occur in two ways: a case can be misclassified as a control or a control can be misclassified as a case. For example, the Prostate Specific Antigen (PSA) test is used to diagnose prostate cancer. Not all men who have prostate cancer show elevated PSA, which results in a case being misclassified as a control (Gordon and Finch, 2005). Lack of consideration of underlying subtypes leads to imprecise phenotyping. Prior work has shown that imprecise phenotyping significantly reduces the effect sizes for models of genetic association and contributes to the missing heritability problem (Sluis et al., 2010).

One way to address phenotypic heterogeneity is disease subtyping using unsupervised statistical techniques with phenotypic information such as symptoms or using one or multiple types of omics data. Existing approaches for disease subtyping fall roughly into three categories. Approaches in the first category use phenotypic data and are highly applicable to psychiatric disorders. Various clustering processes such as Latent Class Analysis (LCA) (Grados et al., 2008; Prust et al., 2011) and finite mixture models (Wessman et al., 2009) are used based on secondary phenotypic measures on disease cases. The second category of approaches utilize one type of genomic data, such as gene expression or epigenetic data, and are applicable to a wide variety of diseases (Bair and Tibshirani, 2004; Pyatnitskiy et al., 2014). These approaches utilize clustering algorithms and variable selection techniques. The third category of approaches uses multiple types of genomic data, such as gene expression, copy number variation, and DNA methylation. Various statistical tools are used, including models implemented in a Bayesian framework as well as clustering techniques. Examples of these techniques include Shen et al. (2012) and Kristensen et al. (2014). None of the techniques incorporate heterogeneity into an association testing framework; however, doing so, could improve effect size estimation and provide inferences on heterogeneity.

The finite mixture model is a classical statistical technique to model population heterogeneity. Since the work of Pearson (1894), the finite mixture model has been widely used as a flexible method of modeling heterogeneity in fields of study such as medicine, genetics, biology, astronomy, and social sciences. Many well-developed statistical techniques in these fields, e.g., latent class analysis, cluster analysis, discriminant analysis, image analysis, and survival analysis, are underpinned by finite mixture model theory. The finite mixture model is ideal in situations where covariates that characterize the distribution of the variable under consideration are missing because they were not observed or measured during the data collection process. Typically, the finite mixture model is used to model data that arises from a heterogeneous population. When the population can be divided into some number of sub-populations, and for each sub-population the data can be modeled by a parametric distribution, then their mixture gives the marginal distribution for the population. Here we implement a finite mixture of logistic regression models to incorporate heterogeneity into an association framework. Simulation studies are used to show that the effect size estimates have lower bias compared to classical logistic regression while maintaining comparable power.

## 1 Methods

### 1.1 Finite Mixture Model and EM algorithm

Let *Y*_1_, *…, Y*_*n*_ be a random sample of size n with observed values *y*_1_, *…, y*_*n*_. Random variable *Y*_*i*_ arises from a finite mixture distribution if the density function can be written in the form,

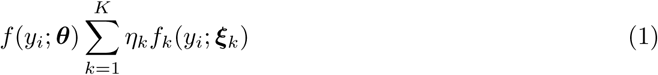

where *f*_*k*_(*y*_*i*_; ***ξ***_*k*_) are parametric density functions with unknown parameters ***ξ***_*i*_, *η*_*k*_ satisfy 0 ≤ *η*_*k*_ ≤ 1 for all *k* with 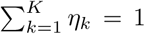, and θ = (*η*_1_, …, *η*_*k*_, ψ^*T*^)^*T*^ where the vector Ψ contains all the parameters in ξ1, …, ξk assumed a priori to be distinct. The quantities *η*_*k*_ are called mixing proportions, and *f*_*k*_(*y*_*i*_; ***ξ***_*k*_) are component densities. The expectation maximization (EM) algorithm is an iterative process to approximate the maximum likelihood estimates of the model parameters that depends on unobserved variables. Each iteration of the EM algorithm consists of 2 steps: the expectation (E) step and the maximization (M) step. In the E-step a function is created for the expectation of the log-likelihood evaluated at the current estimates of the parameters, and in the M-step the parameters are estimated by maximizing the function from the E-step. Consider the mixture model (1). In the finite mixture model the observed data ***y*** = (*y*_1_, *…, y*_*n*_)^*T*^ is viewed as being incomplete as the vectors of component-labels, ***z***_1_, *…*, ***z***_*n*_, are missing, where ***z***_*i*_ is a *K*-dimensional vector with *z*_*ki*_ = 1 or 0, according to whether *y*_*i*_ did or did not arise from the *k*th component. The vectors ***z***_1_, *…*, ***z***_*n*_ are taken to be the realized values of multinomial random variables, ***Z***_1_, *…*, ***Z***_*n*_. The log-likelihood of the complete data, ***y***_*c*_ = (***y***^*T*^, ***z***^*T*^), is given by

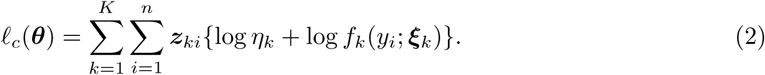

The EM algorithm based on the complete data log-likelihood (2) is described in (Aitkin and Wilson, 1980) and also available in most finite mixture model textbooks such as McLachlan and Peel (2004). A general schemetic of the EM algorithm is given in Figure 1. There are several different methods for choosing the initial values of the parameters as discussed in Karlis and Xekalaki (2003). The stopping criteria are generally based on the difference in likelihood between two consecutive iterations or the Aitken acceleration (McLachlan and Peel, 2004).

**Figure 1:**
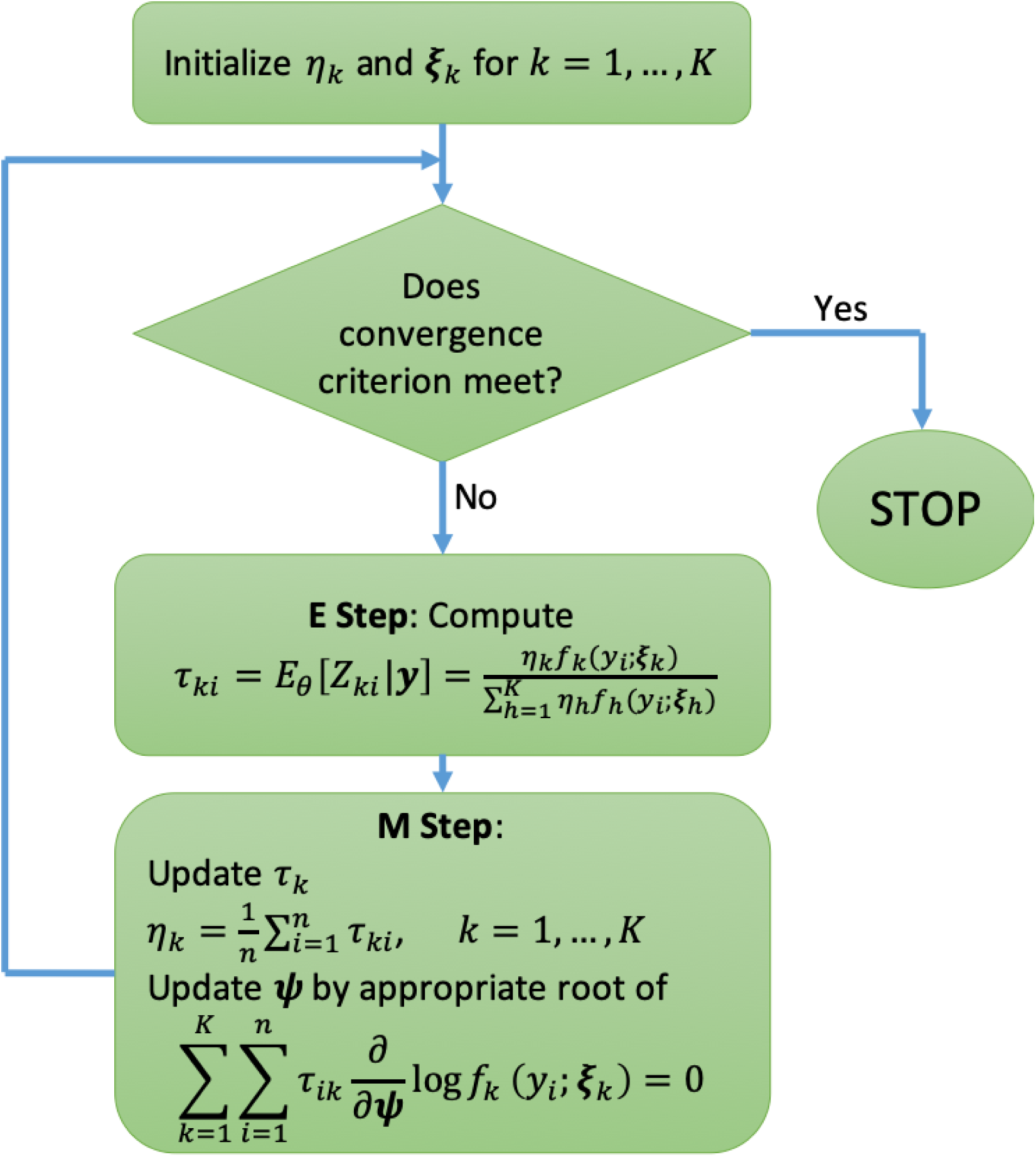
EM algorithm for a finite mixture model.

### 1.2 Bernoulli Mixture Model

Let *Y*_*i*_ and *G*_*i*_ be the disease status and minor allele count (0, 1, or 2) at a bi-allelic genetic locus under consideration for the *i*th observation where, 1 ≤ *I* ≤ *n*. Under an additive model, the logistic regression model is given by,

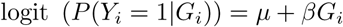

where, *µ* and *β* are regression coefficients. Another form of the model equation is,

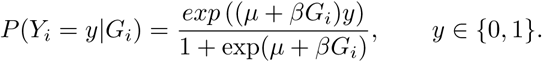

The coefficient *β* is known as the effect size of the locus. Under the hypothesis that the genetic variant is associated with disease for only a part of the population, the disease outcome can be modeled as a finite mixture distribution. We model the distribution of *Y*_*i*_ as a mixture of two component distributions:

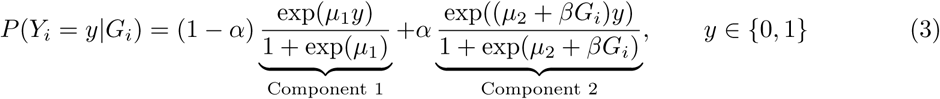

where, *µ*_1_, *µ*_2_, and *β* are regression coefficients and *α* is the mixing parameter that represents the proportion of the population for which the genetic variant is associated with disease. Component 1 and component 2 correspond to the partitions of the population at which the genetic variant has zero and non-zero effect, respectively. Model (3) is a mixture of a generalized linear model. Not all such models are identifiable. For models that lack identifiability, the estimation procedure may not be well-defined and asymptotic theory may not hold (McLachlan et al., 2019). Note that, for model (3), we can establish only one equation which has four unknown parameters:

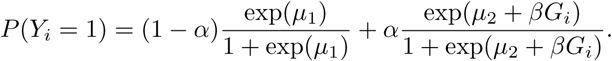

Hence model (3) is not identifiable since different values of the parameters *µ*_1_, *µ*_2_, *α*, and *β* result in the same probability distribution of *Y*. The identifiability problem can be resolved by considering ancestry clusters and modeling the number of diseased individuals in an ancestry cluster to be a binomial random variable. The R package “flexmix” (Grün and Leisch, 2007; Leisch, 2004) is readily available to fit a mixture of generalized linear regression such as the Bernoulli mixture model. In our main model, discussed in following section, we could not implement the package because of the unidentifiability of the model for our data.

### 1.3 Binomial Mixture Model

Identifiability of a mixture of binomial distributions has been well studied. Follmann and Lambert (1991) studied the mixture of binomial regression models, but their model only allows the intercept to follow a mixture distribution, keeping the coefficients of the covariates constant over components. Sufficient conditions for identifiability for a more general model is given in Grün and Leisch (2008). One of their three sufficient conditions is *N* ≥ 2*K* − 1, where *N* is the number of trials of the binomial distribution and *K* is the number of components in the mixture distribution. The other two conditions relate to convergence. In the case of a binary disease outcome, the Bernoulli mixture model does not satisfy the first condition. In this section, we will introduce a binomial mixture model that is identifiable.

#### 1.3.1 Model Definition

In a classical GWAS setting, ancestry covariates are often included in the association model to control for population structure. Here, we propose to use ancestry clusters and define the response variable as the number of cases within an ancestry cluster to solve the identifiability problem. Table 1 shows the data structure after collapsing case-control data for each unique combination of ancestry cluster and genotype. For each ancestry cluster, we have up to three possible genotypes. An instance in the data corresponds to a genotypic value (coded as minor allele count) at an ancestry cluster. At an instance, let *N* be the number of observations (Bernoulli trials) and *Y* be the number of cases (number of successes). As long as all values of *N* are greater than or equal to 3, we have met the first condition for identifiability of a two-component (*K* = 2) binomial mixture model.

**Table 1:** Data structure exemplar for a mixture of binomial distributions. Column AC and G represent the ancestry cluster and genotype, respectively. Column N gives the total number of observations in a ancestry cluster (AC) with minor allele count (G), which is the number of trials for the binomial distribution, and Y represents the number of cases within ancestry cluster AC with minor allele count G.

| AC | G | N | Y |
| --- | --- | --- | --- |
| 1 | 0 | 5 | 1 |
| 1 | 1 | 6 | 2 |
| 1 | 2 | 3 | 2 |
| 2 | 0 | 10 | 1 |
| 2 | 1 | 5 | 2 |
| ⋮ | ⋮ | ⋮ | ⋮ |

Assume *m* ancestry clusters, *i* = 1, 2, *…, m*, and within cluster *i* there are *m*_*i*_ genotype values observed with *m*_*i*_ ∈ {1, 2, 3}. Let *N*_*ij*_, *Y*_*ij*_, and *G*_*ij*_ represent the total number of observations, number of cases, and genotype corresponding to the *i*th ancestry cluster and the *j*th genotype value. The two-component mixture of binomial distributions is given by

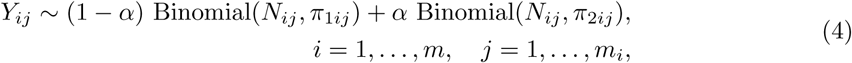

where the probabilities *π*_1*ij*_ and *π*_2*ij*_ are from logistic regression models,

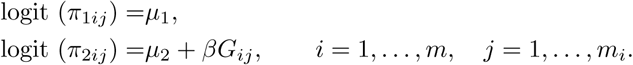

The first component, say *C*_1_, in the mixture model (4) corresponds to the part of the population for which the locus is not associated with disease and the second component, say *C*_2_, corresponds to the associated group. We use an expectation maximization (EM) algorithm to estimate the model parameters.

### 1.3.2 Parameter Estimation

The conditional densities of *Y*_*ij*_ given components *C*_1_ and *C*_2_ are,

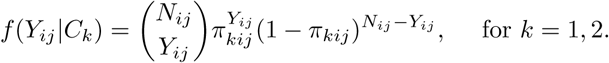

The missing data is a two-dimensional latent class vector ***z***_*ij*_ = (*z*_1*ij*_, *z*_2*ij*_), *i* = 1, *…, m* and *j* = 1, *…, m*_*i*_, defined as,

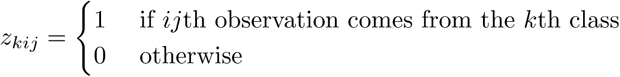

for *k* = 1, 2. The component-label vectors ***z***_*ij*_ are taken to be realizations of the Bernoulli random vectors ***Z***_*ij*_ for *i* = 1, *…, m, j* = 1, *…, m*_*i*_ with density function,

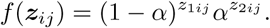

The set of parameters to be estimated is ***θ*** = {*α, β, µ*_1_, *µ*_2_}. The complete data likelihood is given by,

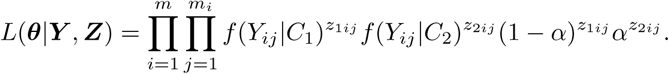

where ***Y*** = (*Y*_*ij*_) and ***Z*** = (*z*_*kij*_), *k* = 1, 2, *i* = 1, *…, m*, and *j* = 1, *… m*_*i*_.

The log likelihood is

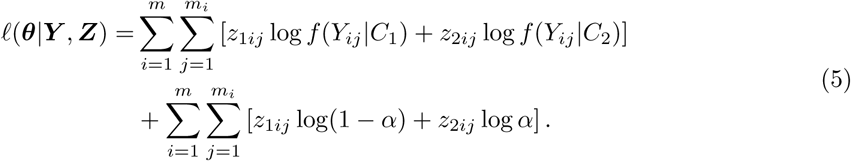

In the expectation step of the EM algorithm we find the conditional expectation *E* [***Z***_*kij*_|***Y***, ***θ***].

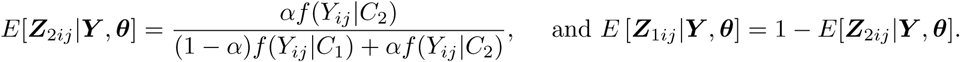

In the maximization step, the log-likelihood (5) is maximized for *α* at:

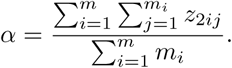

To maximize (5) for *µ*_1_, *µ*_2_, and *β*, note that the second term does not depends on these parameters. We can simplify the first term as follows:

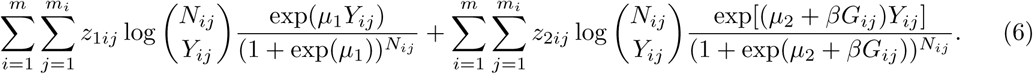

Maximizing (6) is equivalent to maximizing the two terms separately, which can be achieved by fitting separate logistic regression models with weights of *z*_1*ij*_, and *z*_2*ij*_, respectively.

### 1.3.3 Hypothesis Test

We are interested in whether 1) there is phenotypic heterogeneity, and 2) under the assumption of phenotypic heterogeneity, a genetic variant is associated with disease status. Consider testing the null hypothesis: *H*_0_ : *α* = 1 and *β* = 0. Under this null hypothesis, the mixture model (4) reduces to the logistic regression, logit (*π*_1*ij*_) = *µ*_1_. The corresponding alternative hypothesis is *H*_*a*_ : *α ≠* 1 and/or *β ≠* 0. Under this alternative hypothesis we cannot conclude association as there can be a scenario where heterogeneity (*α ≠* 1) exists in terms of a genetic variant, which is not associated (*β* = 0), due to a difference in disease prevalence between two groups in the population (*µ*_1_ *≠ µ*_2_). A more sensible hypothesis can be formed under the assumption that the population prevalence is equal in the two groups, which is a reasonable assumption for a polygenic disease and a genetic variant whose effect size is not extreme. Under the assumption that, *µ*_1_ = *µ*_2_ = *µ* the binomial mixture model becomes:

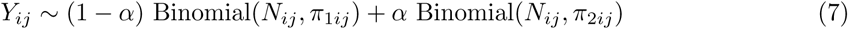

where the probabilities *π*_1*ij*_ and *π*_2*ij*_ are from logistic regression models

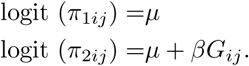

Now we can test the null hypothesis, *H*_0_ : *α* = 1 or *β* = 0, against the alternative, *H*_*a*_ : *α ≠* 1 and *β ≠* 0. Under this null hypothesis, the mixture (7) reduces to a logistic regression: logit (*π*_1*ij*_) = *µ*. Under the alternative hypothesis, we are now able to conclude association of the genetic variant with the trait.

The likelihood ratio test statistic is given by,

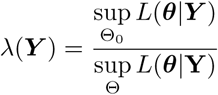

where Θ = [0, 1] × ℝ^3^ is the full parameter space and Θ_0_ is a subset corresponding to the null hypothesis. Under some regularity conditions, −2 log *λ* follows a chi-squared distribution with degrees of freedom equal to the difference in the number of parameters between the full and null model Casella and Berger (2002). The regularity condition violated in model (7) is that the true parameter value, ***θ***_0_, is an interior point in the parameter space. Self and Liang (1987) provides the asymptotic distribution of −2 log *λ* under such non-standard conditions. According to Self and Liang (1987) the LRTS has an asymptotic 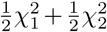 distribution. A Q-Q plot of the LRTS is given in Figure 6.

**Figure 2:**
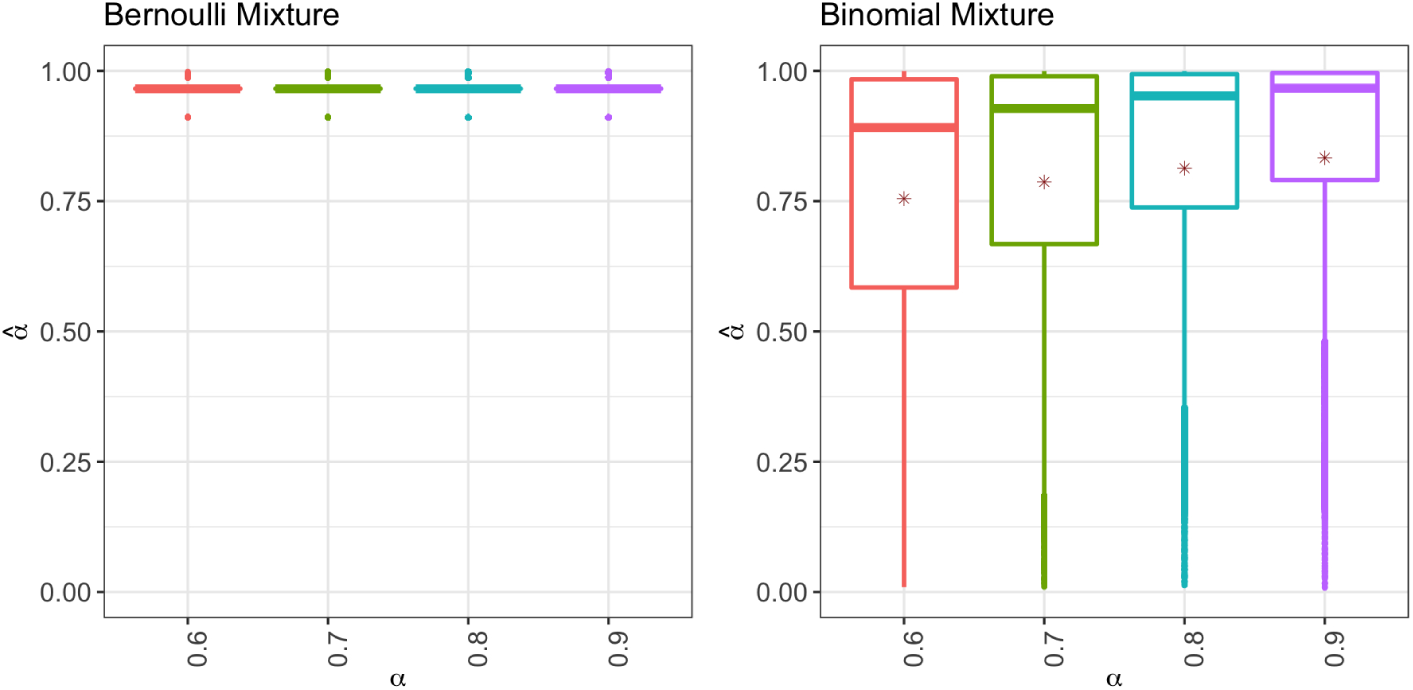
Estimation of the mixing parameter using a Bernoulli mixture model and a binomial mixture model. The *x* and *y* axes show the mixing parameter, *α*, and corresponding estimates,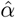, respectively. The estimates correspond to simulations with all possible combinations of maf and *β*. The * on the boxplots represents the mean of 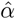.

**Figure 3:**
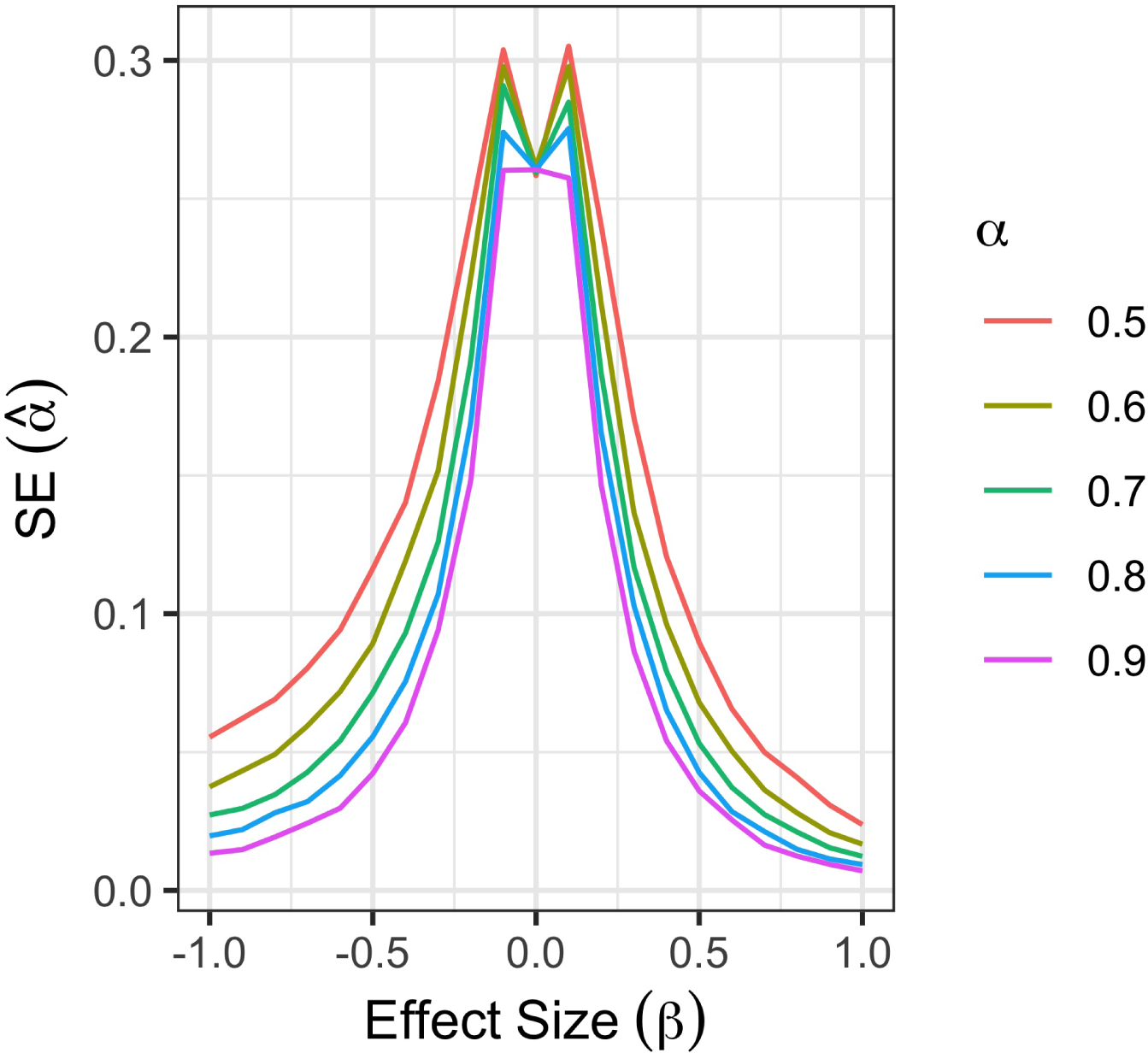
Standard error of mixing proportion estimates 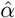 Results are based on simulations with disease prevalence *K* = 0.02 and combined over different levels of allele frequencies. The horizontal axis represents the effect size used in simulations, and the vertical axis represents the standard error of 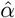 calculated from 1,000 replicates.

**Figure 4:**
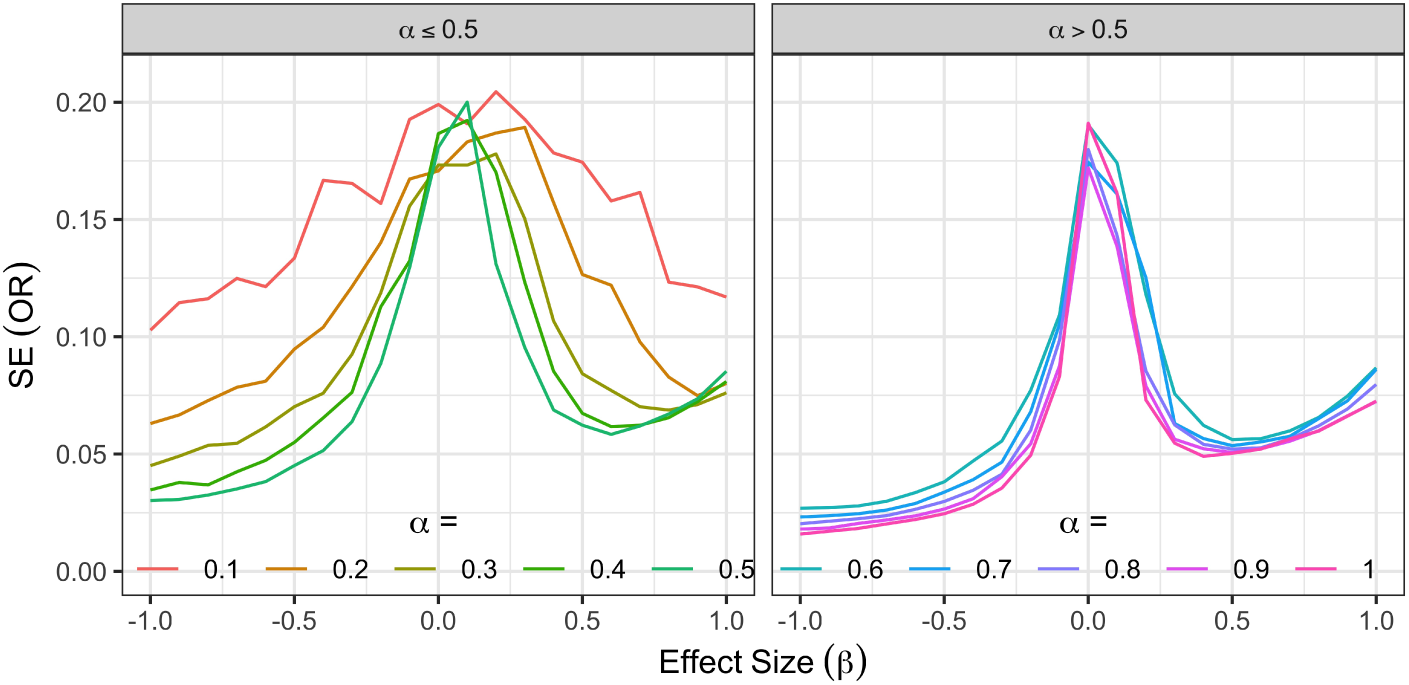
Standard error of odds ratio estimates. The horizontal and vertical axes have the OR and SE of estimated OR, respectively. The left and right panes show results for *α ≤* 0.5 and *α >* 0.5, respectively.

**Figure 5:**
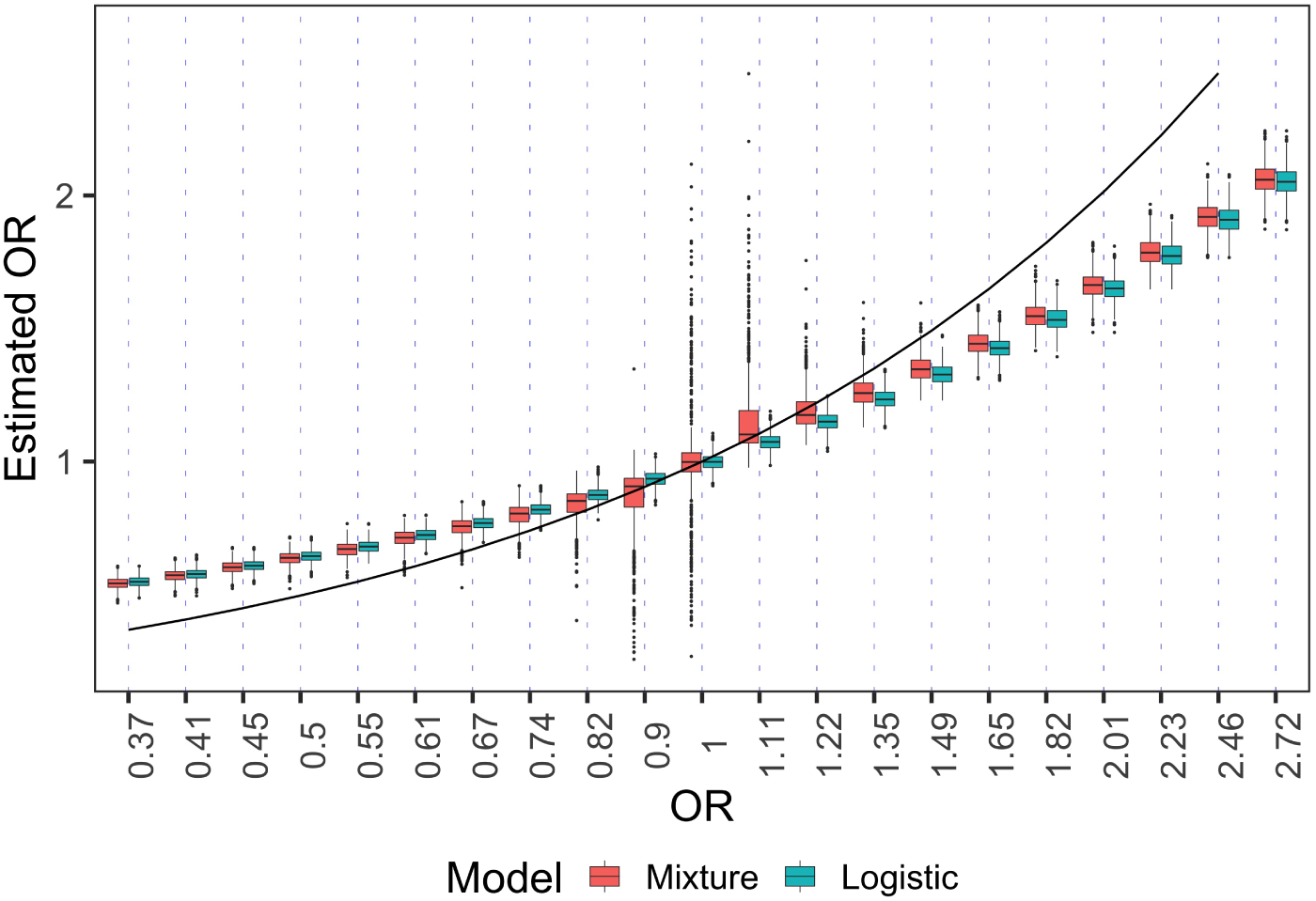
Comparison of the binomial mixture model and the logistic regression in terms of bias in odds ratio estimation. A value closer to the black curve yields smaller bias. The results are based on 1,000 samples with 5,000 cases and 20,000 controls, each considering disease prevalence of 2%, maf of 0.15 and proportion of associated group 70%.

**Figure 6:**
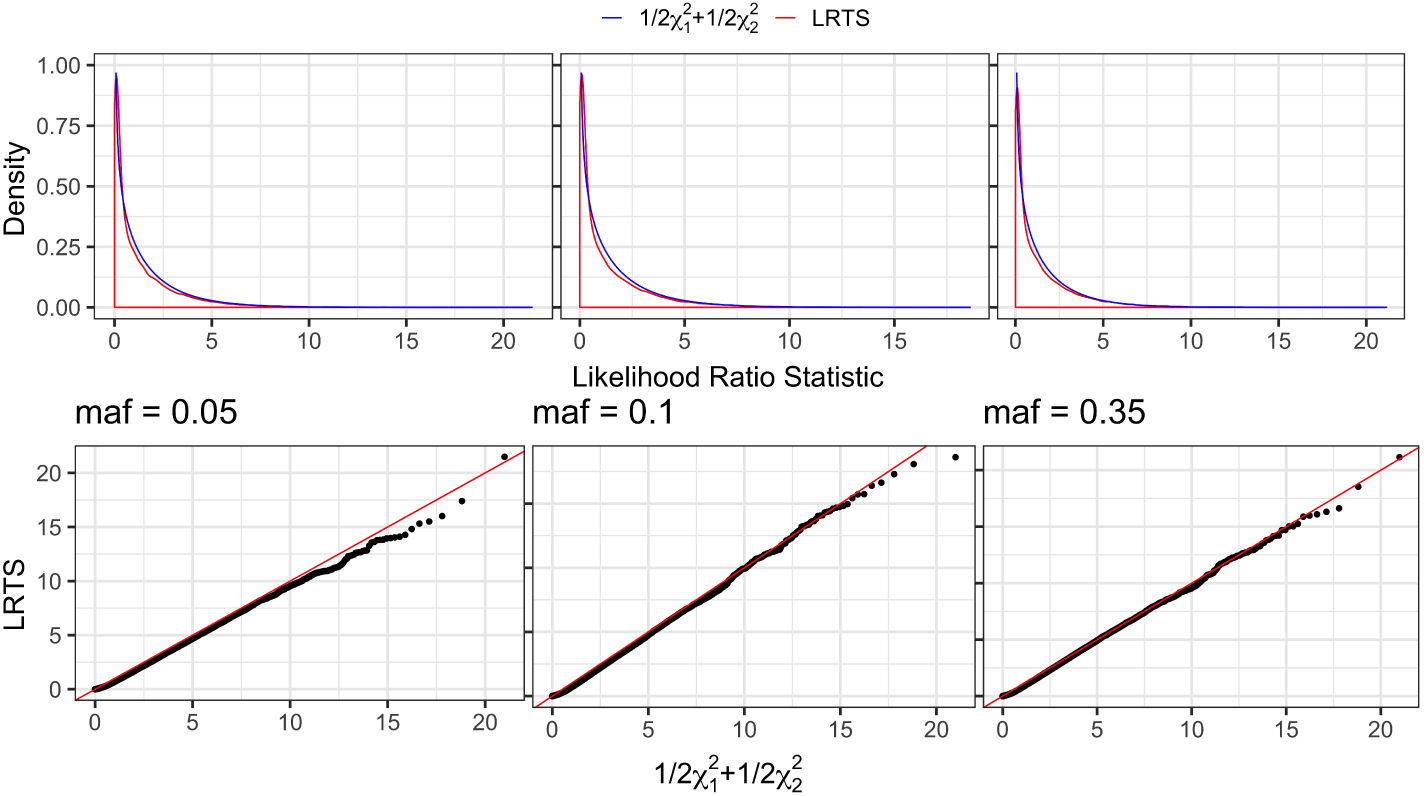
Under the null hypothesis, the likelihood ratio test statistic has an asymptotic 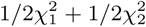 distribution. The top panel of the figures show density plots of the LRTS and the bottom panel shows Q-Q plots. Three columns (from left to right) represent maf of 0.10, 0.25, and 0.40. For each allele frequency, the results are based on 31,000 simulated samples with *K* = 0.02, and either *β* = 0 or *α* = 0.

### 1.4 Simulation

#### 1.4.1 Simulation Model

Assume that we want to simulate *N*_1_ cases and *N*_2_ controls from a population that is divided into two subgroups *S*_1_ and *S*_2_ of proportions 1 − *α* and *α*, respectively. The subgroup *S*_2_ represents the group of individuals for which the genetic variant is associated with disease status, and *S*_1_ represents the non-associated group. The subgroups are considered at the population level, so they contain both cases and controls. Let the effect size in the association group be *β*. Let *p*_1_, *p*_2_, *K*_1_, and *K*_2_ be the minor allele frequencies (maf) and disease prevalences for *S*_1_ and *S*_2_, respectively. The number of cases *N*_11_ drawn from *S*_1_ follows a Binomial (*N*_1_, 1 − *α*) and so we have *N*_12_ = *N*_1_ − *N*_11_ cases from *S*_2_. Similarly, we have *N*_21_ ∼ Binomial (*N*_2_, 1 − *α*) and *N*_22_ = *N*_2_ − *N*_21_ controls from *S*_1_ and *S*_2_, respectively.

Observations were randomized to one of the *m* ancestry clusters. We expect that the allele frequencies will differ among the clusters from the overall allele frequencies *p*_1_ and *p*_2_. Let *F*_*ST*_ be the fixation index. Between-population variation in allele proportions at a biallelic locus is often modeled by the Beta distribution with parameters *λp* and *λ*(1 − *p*), where 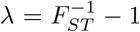 (Balding and Nichols, 1995). We will denote the maf at the *i*th ancestry cluster by 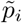.

The disease status does not depend on genotype in the non-associated group, so the genotype distribution of cases and controls are obtained assuming Hardy-Weinberg equilibrium (HWE). For the cases and controls from *S*_2_, the genotype distribution can be obtained using Bayes theorem for *g* ∈ {0, 1, 2} as

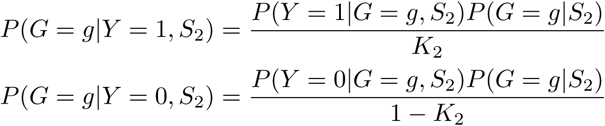

where *P* (*Y* = *y*|*G* = *g, S*_2_) is obtained from a logistic regression model as,

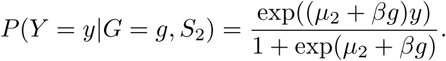

The constant coefficient *µ*_2_, specific to the *i*th ancestry cluster, satisfies the following equation:

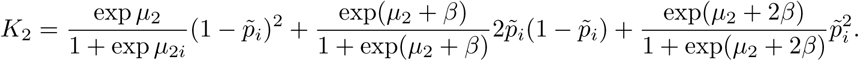

#### 1.4.2 Simulation Parameters

Diseases with four different levels of prevalence were simulated: 0.01, 0.02, 0.05, and 0.1. The overall population prevalence is decomposed into prevalences for the two sub-groups, *S*_1_, and *S*_2_ as,

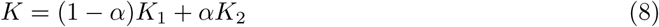

For a complex disease, one single genetic variant is not expected to have a large effect on disease prevalence. If a variant is deleterious, we can expect a higher prevalence in the *S*_2_ group than *S*_1_. On the other hand, if a genetic variant is beneficial, we can expect a lower prevalence in *S*_2_ compared to *S*_1_. For simulation, we set *K*_2_ to be 5% higher or lower than *K* and calculate *K*_1_ accordingly from (8).

We simulate genetic variants with maf from 5% to 50% with an increment of 5%. Since it is highly unlikely that allele frequency at a particular locus is driven by the fact that the locus is associated with a complex trait, we assume that the allele frequency in *S*_1_ and *S*_2_ are equal.

In our simulation, we consider the mixing parameter *α* to vary from 0.1 to 1 with an increment of 0.1. The effect size *β* is taken to fall in a range of -1 to 1 in increments of 0.1. Theoretically, the fixation index, *F*_*ST*_ varies from 0 to 1, but it is rare to have *F*_*ST*_ more than 0.3. In most human populations, such as in Europeans, the fixation index is reported to be about 0.01 (Tian et al., 2009) which is used in our simulation. For each point in the parameter space consisting of *K, p, α*, and *β*, we performed 1,000 simulations with 5,000 cases and 20,000 controls.

## 2 Results

### 2.1 Use of Binomial Mixture to Meet Identifiability Conditions

We have discussed the identifiability issue of the Bernoulli mixture model (3) that motivated us to develop the binomial mixture model (7). Figure 2 shows the impact of non-identifiability of the estimates of the mixing proportion based on simulations with 2,000 cases and 8,000 controls. If we use the Bernoulli mixture model in equation (3), almost all observations are assigned to the association group, *S*_2_, whereas the mean of the mixing parameter estimates are comparatively closer to the truth for the binomial mixture model in equation (7). For example, for *α* = 0.6, 0.7, 0.8, and 0.9, the mean estimates are 0.67, 0.72, 0.77, and 0.85, respectively. The estimates are upward biased and lack precision. For example, the standard deviation of 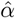 at *α* = 0.7 is 0.29. As shown in Figure 3, the standard error of the mixing proportion estimate decreases as the magnitude of the effect size increases. For a genetic variant with moderate to high effect (*β* ≥ 0.5), 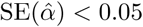 when mixing proportion *α* ≥ 0.7. For a fixed effect size, a higher mixing proportion is estimated with more precision, i.e., smaller 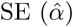.

### 2.2 OR Estimates are More Precise for Higher Values of OR and Mixing Proportion

Figure 4 shows the standard error of the odds ratio estimates for *α* = 0.1,0.2,*…*, 0.5 against the effect size, *β*. The standard error of the odds ratio estimates decreases with an increase in mixing proportion and OR. With a small mixing proportion, the effective number of cases, i.e., the cases in the associated group, is small. For example, with *α* = 0.1 the effective number of cases is only 500. Such a small sample size does not provide precise estimates of effect sizes. In the extreme case of *α* = 0.1 the standard error varies between 0.1 and 0.2. Given this lack of precision, only results for simulations with mixing proportion higher than 0.50 will be provided.

For a protective genetic variant (OR *<* 1), the standard error of the OR estimate decreases with a decrease in OR. For a deleterious genetic variant (OR *>*1), the standard error of the OR estimate decreases from low to moderate effect sizes and increases slightly for higher effect sizes (Figure 4). For a fixed OR, the standard error of the OR estimate is smaller for a higher level of *α*.

### 2.3 Binomial Mixture Model Yields Smaller Bias Compared to Logistic Regression

For all simulations with mixing proportions from 0.60 to 0.90, for all levels of maf, disease prevalence, and positive *β*, the odds ratio estimates using the logistic regression and the binomial mixture have downward bias. The bias is upward for negative *β*. The binomial mixture model reduces the odds ratio bias compared to the logistic regression. Table 2 shows the percentage of bias reduced using the binomial mixture model compared to logistic regression for the simulations with *K* = 0.02. The percentages are calculated as 100 × (Bias_Logistic – Bias_(Binomial Mixture)) / Bias_Logistic. For simulations with *K* = 0.02, the bias reduction by binomial mixture model varies from 1% to 410%. Large bias reduction (*>* 100%) corresponds to small effects where both models have very small bias (Figure 5). Small bias reduction occurs for a high effect genetic variant where both of the models have high bias. For a complex trait, most of the associated genetic variants have moderate effect sizes (0.3 ≤ *β* ≤≤ 0.7). For a genetic variant with moderate effect, the binomial mixture reduces bias in the range of 2.32% to 40.76%. We see similar results for protective loci (*β <* 0) and for all levels of disease prevalence considered. A similar tables for *K*=0.01, 0.05, 01.0 are contained in Supplementary Materials. A paired-sample t-test was performed on the bias of the odds ratio based on simulations with *K* = 0.02, mixing proportion 0.60 to 0.90, and all levels of maf considered. In the presence of heterogeneity (0 *< α <* 1), this hypothesis test concludes that the binomial mixture model yields significantly less bias than logistic regression (p-value *<* 2.2×10^*−*16^). The performance of the binomial mixture model in terms of bias reduction is graphically shown in Figure 5. The figure shows the results from 1,000 simulation using *K* = 0.02, maf = 0.15, and mixing proportion, *α* = 0.70.

**Table 2:**
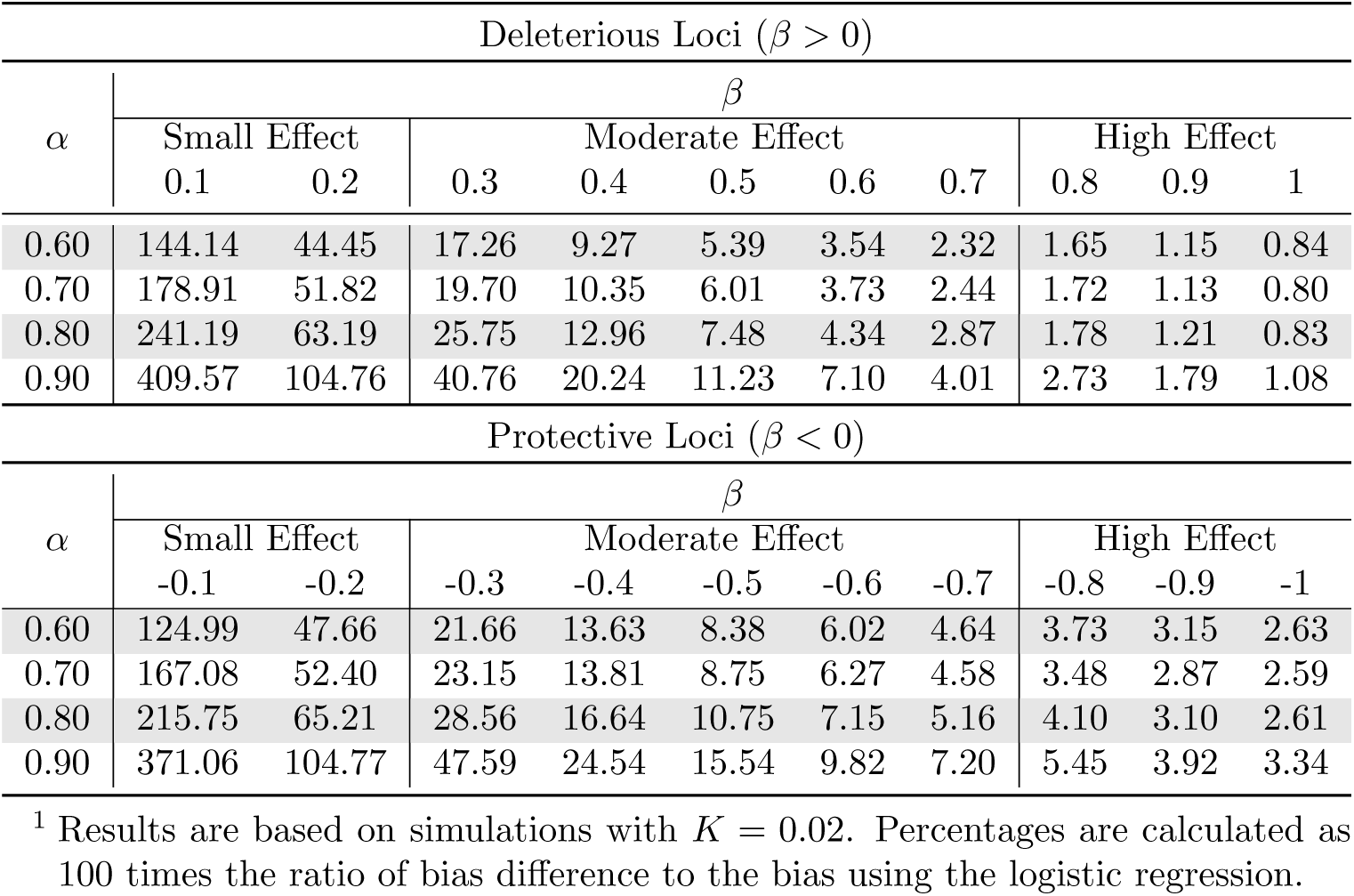
Percentage of bias reduced by the binomial mixture model compared to the logistic regression for simulations with *K* = 0.02.^1^

### 2.4 Establishing the Null Distribution of The Hypothesis Test

There are 1,240,000 simulated sets of data corresponding to the null hypothesis *H*_0_ : *α* = 0 or *β* = 0 over different levels of *K* and maf. The likelihood ratio test statistic (LRTS) did not exceed the critical value of 32.491245 corresponding to a genome-wide significance level (5 × 10^*−*8^) for the distribution of 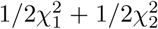 for any set of data. The Q-Q and density plots in Figure 6 shows that the distribution of the LRTS has the same shape as the theoretical distribution.

### 2.5 Power Comparison of the Binomial Mixture Model with Logistic Regression

While the classical hypothesis test in logistic regression framework pertains to association, the hypothesis test in the binomial mixture model provides information about heterogeneity. To compare the two methods, we compared the compound hypothesis of the proposed method with the simple hypothesis of logistic regression. Figure 7 shows the power curves for disease prevalence, *K* = 0.02. Similar figures for *K* = 0.01, 0.05 and 0.1 are given in the supplementary material. When both the allele frequency and mixing proportion are small, none of the models have high power to detect association because of small effective sample size. Power increases with allele frequency and mixing proportion. When the mixing proportion is close to one, the models have comparable power; otherwise, the logistic regression is slightly more powerful in detecting association.

**Figure 7:**
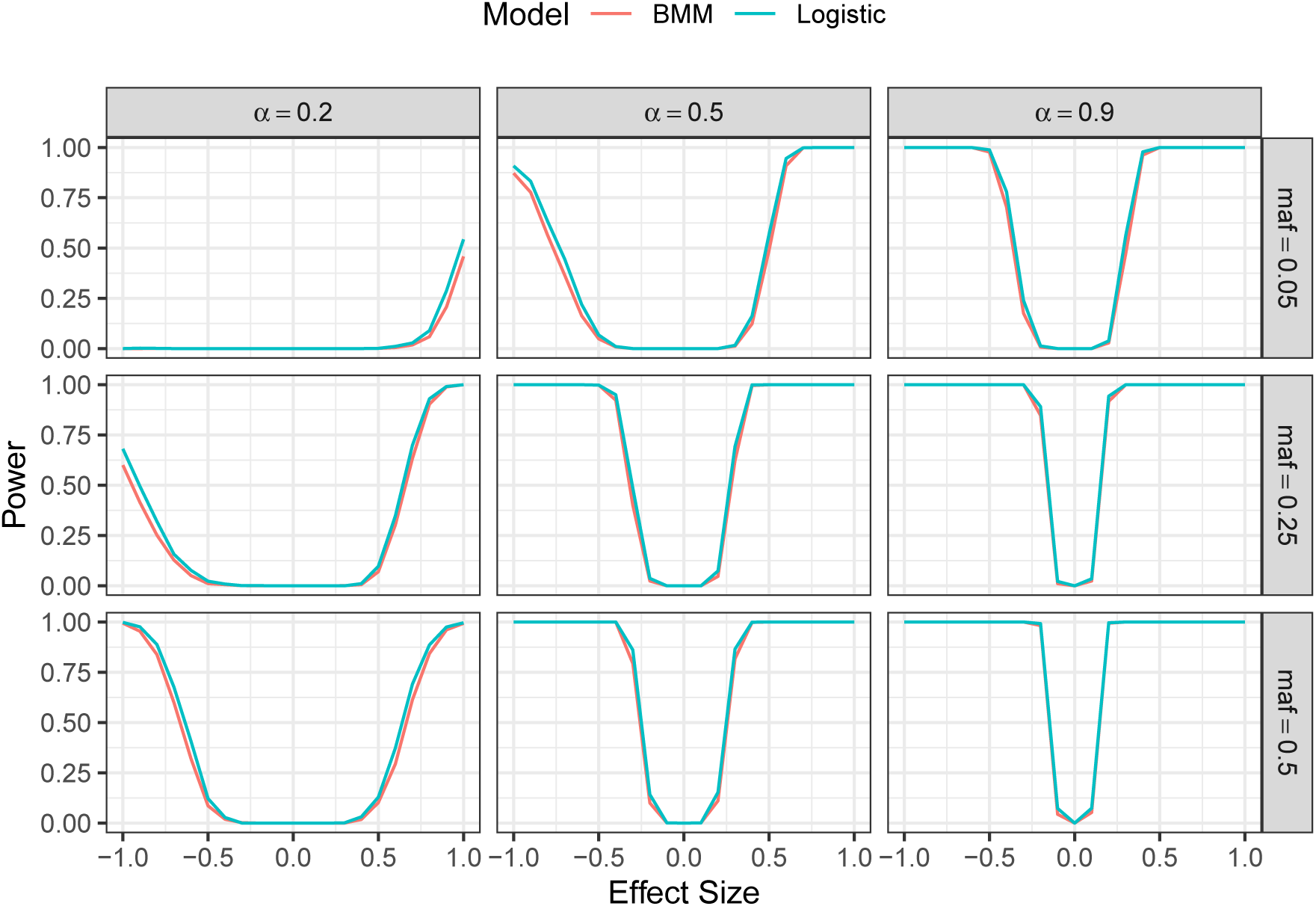
Power curves for hypothesis tests using the proposed mixture model and logistic regression based on samples with disease prevalence, *K* = 0.02. Three rows corresponds to maf of 0.05, 0.25, and 0.50 and three columns corresponds to mixing proportions of 20%, 50% and 90%.

## 3 Discussion

The goal of the study was to incorporate heterogeneity into an association model for a case-control analysis. Finite mixture models are commonly used to model heterogeneity. A finite mixture model for a binary outcome is not identifiable, so is not directly applicable to a case-control study. Utilizing ancestry clusters, we produce an identifiable binomial mixture model. We fit a binomial mixture model to a special form of data. An instance in the data corresponds to a genotype in an ancestry cluster. For identifiability each instance has to have at least 3 observations. For real data analysis, the matched control samples can be selected so that this condition is met.

Compared to logistic regression, the binomial mixture model yields lower odds ratio bias, especially for moderate effect sizes. Performance of the proposed model, in terms of bias reduction, was investigated using simulated data for different levels of disease prevalence, allele frequency of the genetic variant, and proportion of the associated group. In addition to bias reduction, the simulation study shows that the binomial mixture model does not suffers much from power loss compared to logistic regression. A likelihood ratio test was performed to assess the hypotheses: *H*_0_ : *α* = 0 or *β* = 0 versus *H*_*a*_ : *α ≠* 0 and *β ≠* 0. Under the null hypothesis, the LRTS follows a 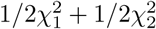 distribution. Using simulated data, we validated the null distribution of the LRTS. We compared power of the compound hypothesis test that pertain to association and heterogeneity with a simple hypothesis test in classical logistic regression that tests for only association. Though the binomial mixture model does not gain power in detecting association, it indicates the presence of heterogeneity and reduces bias.

A common genetic variant only explains a small proportion of the total variability for a complex trait. A precise estimate of the associated group is challenging. Though the estimate of mixing proportion is seen to have high standard error when combined over all simulation scenarios, it is more precise when the proportion of associated group is high, and the genetic variant has higher effect on disease outcome. In the case of a small proportion of the associated group (*α <* 0.5), a larger sample should reduce the variances in estimating the mixing parameter, *α* and effect size, *β*. A precise estimate of the mixing parameter is not expected since subtypes of a complex disease are generally unattainable based on association of a single genetic variant with the disease. Still, even a noisy estimate and indication of heterogeneity may provide important information about the disease architecture.

A more accurate estimate of odds ratios using the binomial mixture could benefit other analyses that uses GWAS summary level data, such as polygenic risk scores (PRS). We expect that the predictive power of PRS would improve using more unbiased estimates of the effect sizes.

The model can be extended to use detailed phenotypes instead of case-control status, and in that way the model would be able to use the underlying structure of a trait. Another possible extension of the binomial mixture model is a multi-SNP model that uses a set of genetic variants from a specific pathway or a gene. Such extensions could lead us to a more precise estimate of the proportion of the associated group. A precise estimate of the mixing proportion should be influential to the drug discovery process since it indicates the proportion of the target population for a specific drug.

## Data Availability Statement

All the codes that were used to simulate the data are available at https://github.com/spaul-genetics/BMM. The github repository also includes all the simulation results.

## A Supplementary Materials

In the supplementary materials we provide additional tables and figures. In the results section we provided a table that describes the percentage of bias reduction using the Binomial Mixture Model based on simulations with disease prevalence, *K* = 0.02. Here we provide similar tables for *K* = 0.01, 0.05 and 0.10. In the tables, *α* represents the mixing proportion, i.e., the proportion of the associated group. The parameter *β* represents the effect size of the genetic variant.

### A.1 Supplementary Tables

**Table A1:**
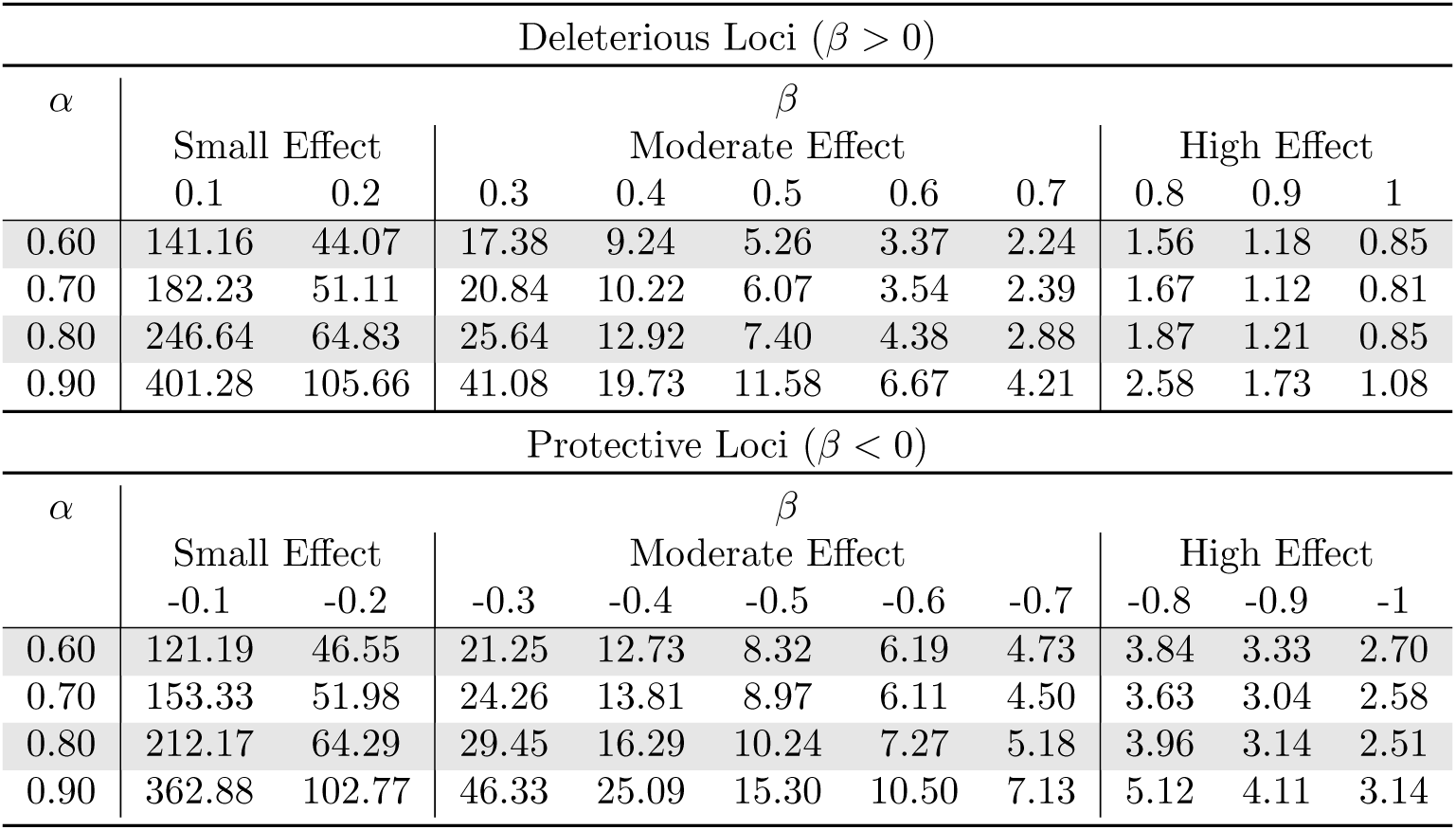
Percentage of bias reduced by the binomial mixture model compared to the logistic regression for simulations with *K* = 0.01. Results are based on simulations with *K* = 0.01. Percentages are calculated as 100 times the ratio of bias difference to the bias using the logistic regression.

| Deleterious Loci ($\beta > 0$) | | | | | | | | | | |
| --- | --- | --- | --- | --- | --- | --- | --- | --- | --- | --- |
| $\alpha$ | $\beta$ | | | | | | | | | |
|  | Small Effect |  | Moderate Effect |  |  |  |  | High Effect |  |  |
|  | 0.1 | 0.2 | 0.3 | 0.4 | 0.5 | 0.6 | 0.7 | 0.8 | 0.9 | 1 |
| 0.60 | 141.16 | 44.07 | 17.38 | 9.24 | 5.26 | 3.37 | 2.24 | 1.56 | 1.18 | 0.85 |
| 0.70 | 182.23 | 51.11 | 20.84 | 10.22 | 6.07 | 3.54 | 2.39 | 1.67 | 1.12 | 0.81 |
| 0.80 | 246.64 | 64.83 | 25.64 | 12.92 | 7.40 | 4.38 | 2.88 | 1.87 | 1.21 | 0.85 |
| 0.90 | 401.28 | 105.66 | 41.08 | 19.73 | 11.58 | 6.67 | 4.21 | 2.58 | 1.73 | 1.08 |
| Protective Loci ($\beta < 0$) | | | | | | | | | | |
| $\alpha$ | $\beta$ | | | | | | | | | |
|  | Small Effect |  | Moderate Effect |  |  |  |  | High Effect |  |  |
|  | -0.1 | -0.2 | -0.3 | -0.4 | -0.5 | -0.6 | -0.7 | -0.8 | -0.9 | -1 |
| 0.60 | 121.19 | 46.55 | 21.25 | 12.73 | 8.32 | 6.19 | 4.73 | 3.84 | 3.33 | 2.70 |
| 0.70 | 153.33 | 51.98 | 24.26 | 13.81 | 8.97 | 6.11 | 4.50 | 3.63 | 3.04 | 2.58 |
| 0.80 | 212.17 | 64.29 | 29.45 | 16.29 | 10.24 | 7.27 | 5.18 | 3.96 | 3.14 | 2.51 |
| 0.90 | 362.88 | 102.77 | 46.33 | 25.09 | 15.30 | 10.50 | 7.13 | 5.12 | 4.11 | 3.14 |

**Table A2:**
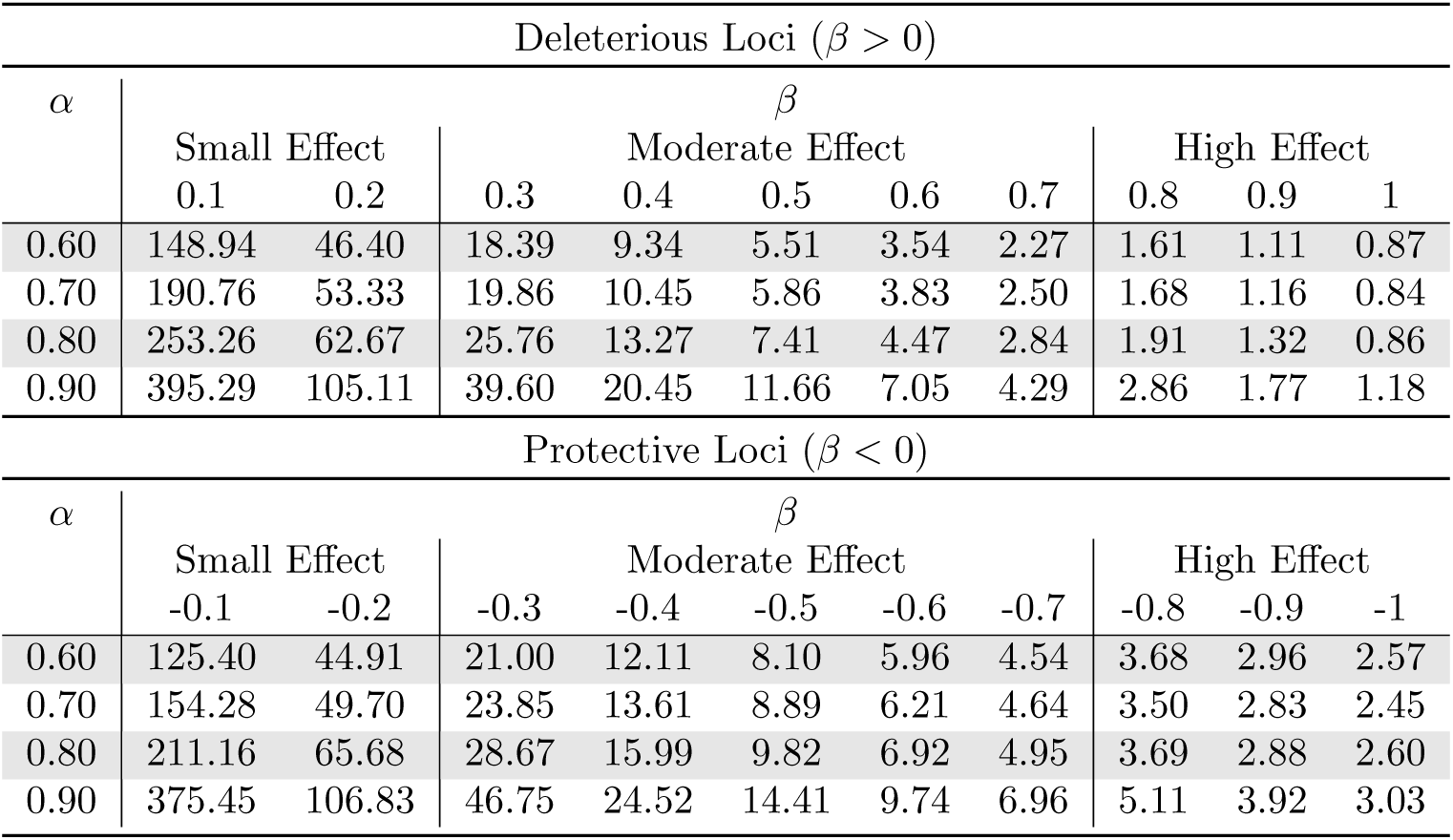
Percentage of bias reduced by the binomial mixture model compared to the logistic regression for simulations with *K* = 0.05. Results are based on simulations with *K* = 0.05. Percentages are calculated as 100 times the ratio of bias difference to the bias using the logistic regression.

**Table A3:** Percentage of bias reduced by the binomial mixture model compared to the logistic regression for simulations with *K* = 0.10. Results are based on simulations with *K* = 0.10. Percentages are calculated as 100 times the ratio of bias difference to the bias using the logistic regression.

| Deleterious Loci ($\beta > 0$) | | | | | | | | | | |
| --- | --- | --- | --- | --- | --- | --- | --- | --- | --- | --- |
| $\alpha$ | $\beta$ | | | | | | | | | |
|  | Small Effect |  | Moderate Effect |  |  |  |  | High Effect |  |  |
|  | 0.1 | 0.2 | 0.3 | 0.4 | 0.5 | 0.6 | 0.7 | 0.8 | 0.9 | 1 |
| 0.60 | 245552.35 | 45.71 | 17.96 | 8.99 | 5.66 | 3.47 | 2.41 | 1.61 | 1.21 | 0.84 |
| 0.70 | 179.78 | 51.58 | 19.66 | 10.68 | 6.17 | 3.63 | 2.51 | 1.79 | 1.23 | 0.89 |
| 0.80 | 486.94 | 64.05 | 24.84 | 12.93 | 7.77 | 4.73 | 3.11 | 1.90 | 1.33 | 0.89 |
| 0.90 | 522.51 | 107.44 | 42.22 | 19.91 | 11.86 | 7.50 | 4.64 | 2.76 | 1.82 | 1.31 |

| Protective Loci ($\beta < 0$) | | | | | | | | | | |
| --- | --- | --- | --- | --- | --- | --- | --- | --- | --- | --- |
| $\alpha$ | $\beta$ | | | | | | | | | |
|  | Small Effect |  | Moderate Effect |  |  |  |  | High Effect |  |  |
|  | -0.1 | -0.2 | -0.3 | -0.4 | -0.5 | -0.6 | -0.7 | -0.8 | -0.9 | -1 |
| 0.60 | 125.32 | 45.44 | 21.84 | 12.73 | 8.40 | 5.89 | 4.43 | 3.43 | 3.03 | 2.43 |
| 0.70 | 157.69 | 51.10 | 24.21 | 13.79 | 8.65 | 5.92 | 4.47 | 3.42 | 2.86 | 2.37 |
| 0.80 | 216.48 | 65.05 | 29.46 | 16.48 | 9.63 | 6.67 | 4.52 | 3.67 | 2.89 | 2.45 |
| 0.90 | 380.55 | 103.07 | 46.88 | 25.44 | 15.10 | 9.89 | 6.99 | 5.12 | 3.59 | 2.89 |

**Figure A1:**
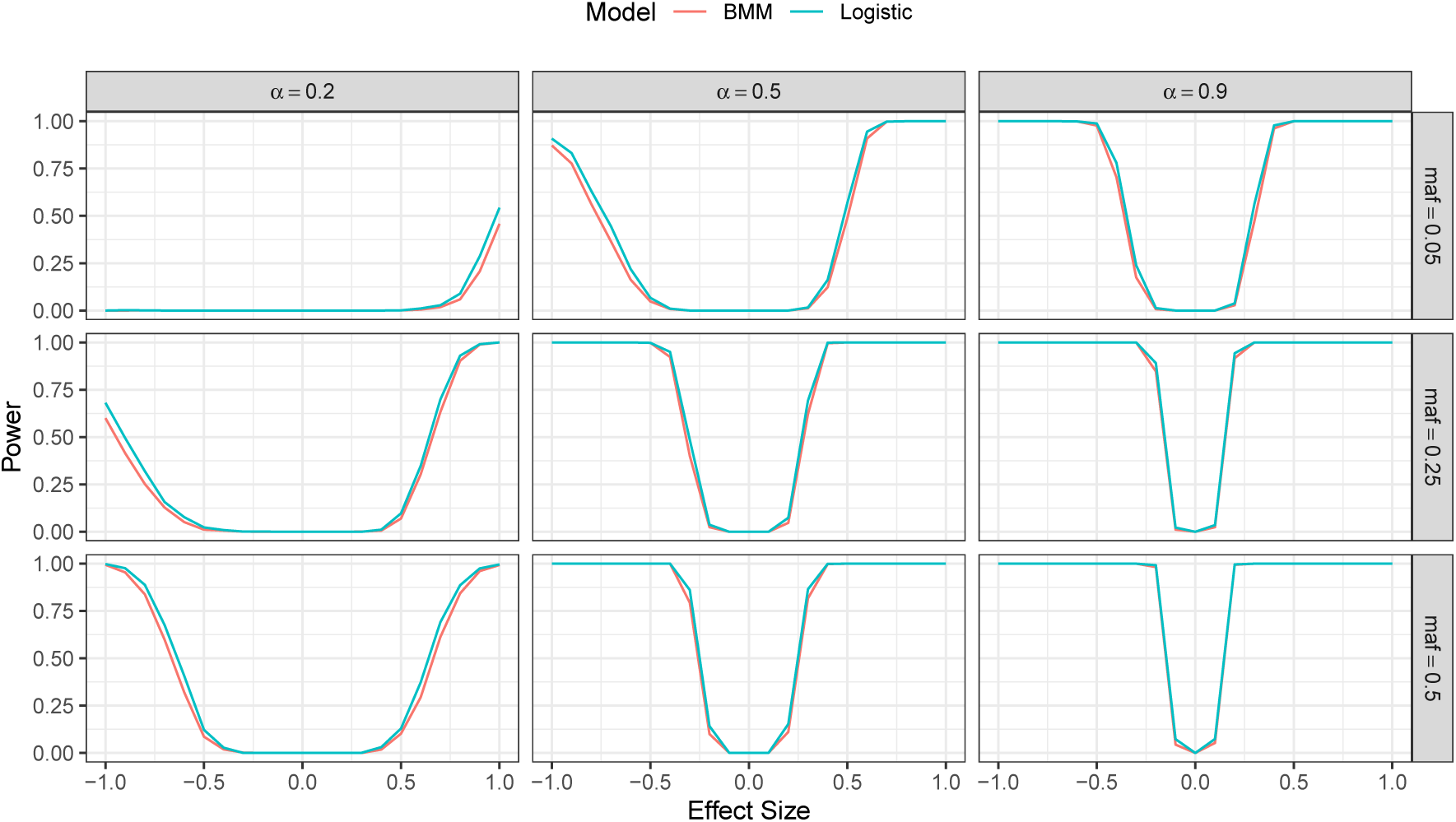
Power curves for hypothesis tests using the proposed mixture model and logistic regression based on samples with disease prevalence, *K* = 0.01. Three rows corresponds to maf of 0.05, 0.25, and 0.50 and three columns corresponds to mixing proportions of 20%, 50% and 90%

**Figure A2:**
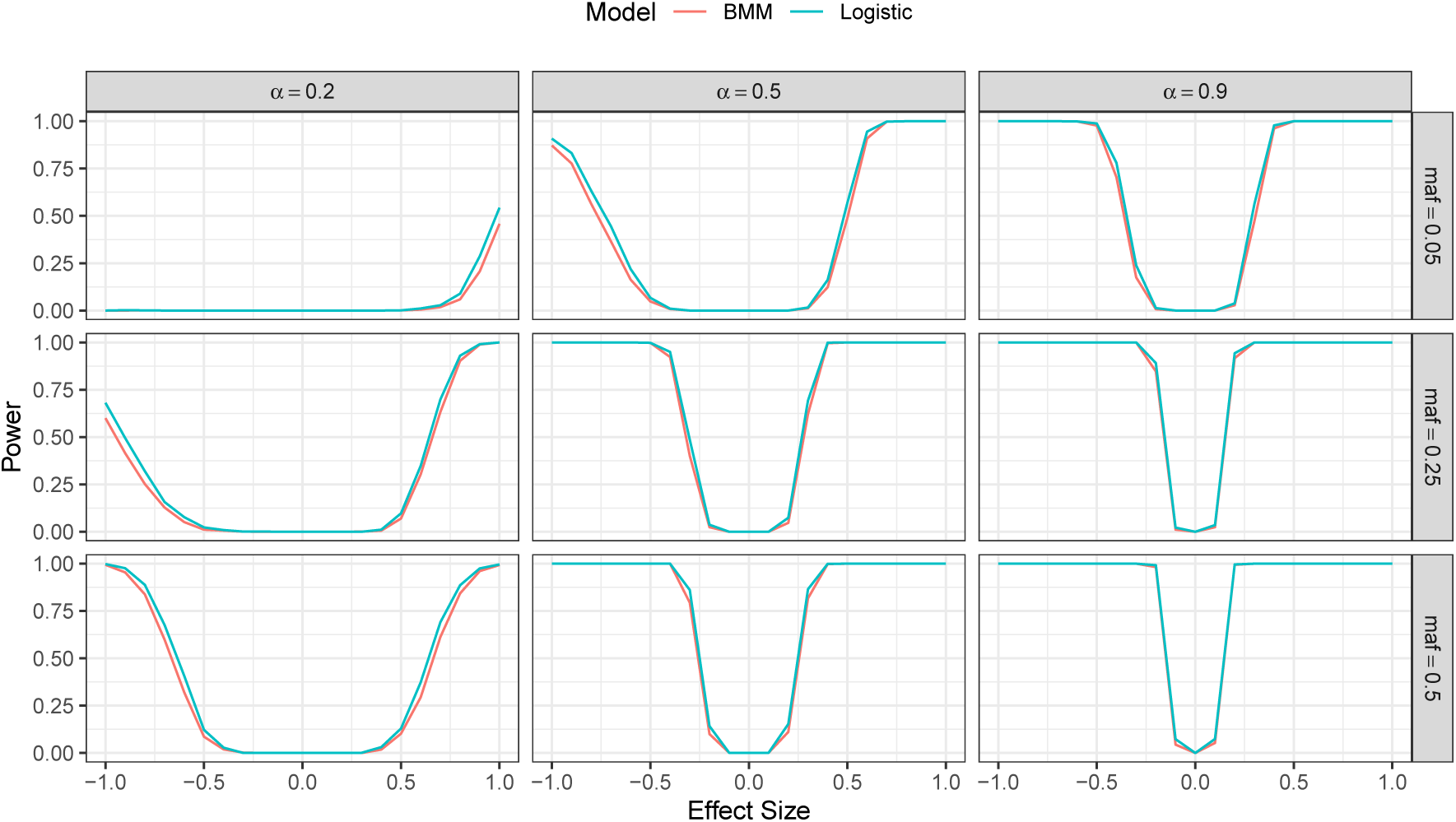
Power curves for hypothesis tests using the proposed mixture model and logistic regression based on samples with disease prevalence, *K* = 0.05. Three rows corresponds to maf of 0.05, 0.25, and 0.50 and three columns corresponds to mixing proportions of 20%, 50% and 90%

**Figure A3:**
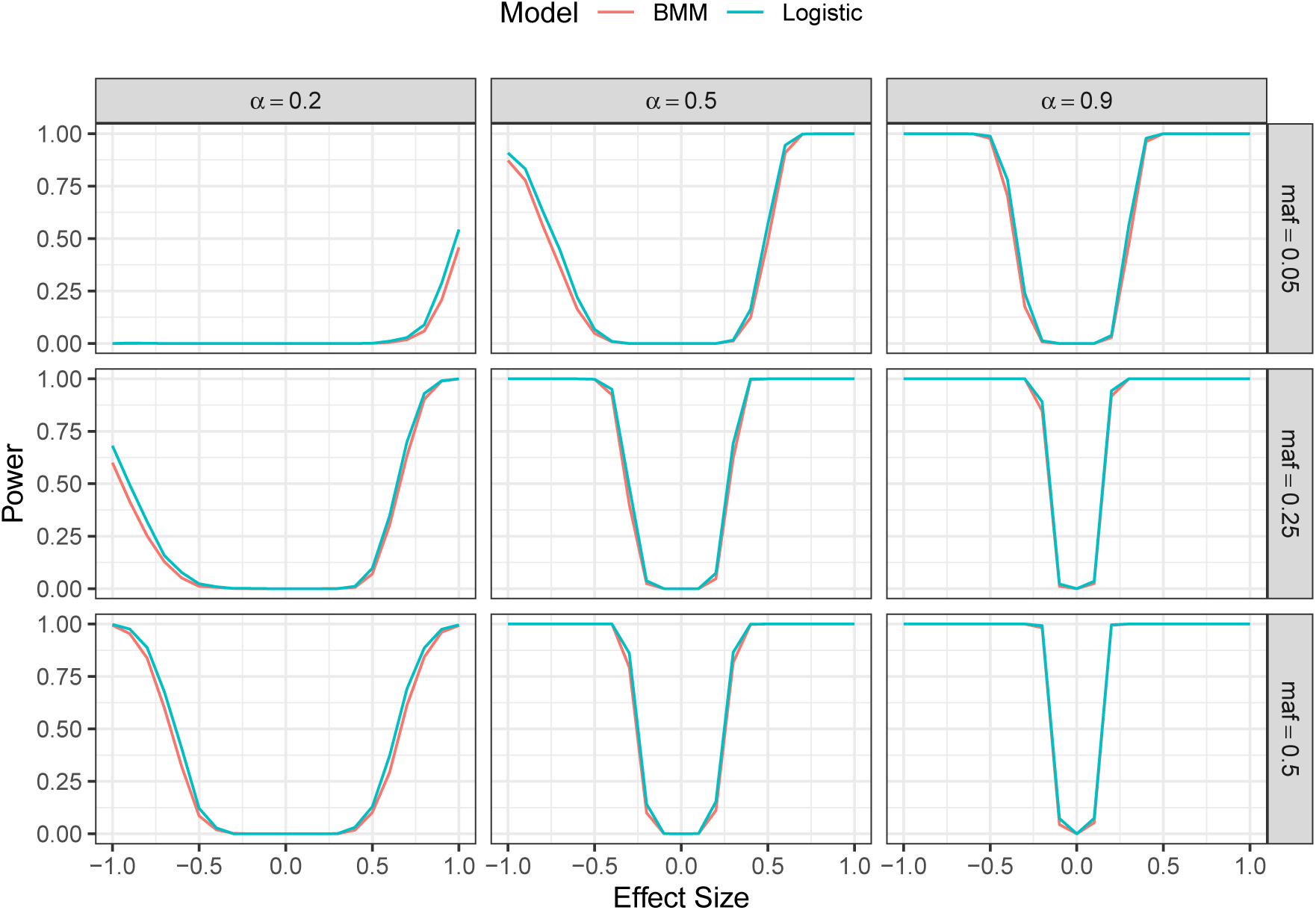
Power curves for hypothesis tests using the proposed mixture model and logistic regression based on samples with disease prevalence, *K* = 0.10. Three rows corresponds to maf of 0.05, 0.25, and 0.50 and three columns corresponds to mixing proportions of 20%, 50% and 90%

